# Stand-alone lipoylated H-protein of the glycine cleavage system enables glycine cleavage and the synthesis of glycine from one-carbon compounds *in vitro*

**DOI:** 10.1101/2021.03.28.437365

**Authors:** Yingying Xu, Yuchen Li, Han Zhang, Jinglei Nie, Jie Ren, Wei Wang, An-Ping Zeng

## Abstract

H-protein, one of the four component proteins (H, T, P and L) of glycine cleavage system (GCS), is generally considered a shuttle protein interacting with the other three GCS-proteins via a lipoyl swinging arm. We report that without P-, T- and L-proteins, lipoylated H-protein (H_lip_) enables GCS reactions in both glycine cleavage and synthesis directions *in vitro*. This apparent catalytic activity is closely related to the cavity on the H-protein surface where the lipoyl arm is attached. Heating or mutation of selected residues in the cavity destroys or reduces the stand-alone activity of H_lip_, which can be restored by adding the other three GCS-proteins. Systematic study of the H_lip_-catalyzed overall GCS reactions and the individual reaction steps provides a first step towards understanding the stand-alone function of H_lip_. The results in this work provide some inspiration for further understanding the mechanism of the GCS and give some interesting implications on the evolution of the GCS.

**Significance statement:** Glycine cleavage system (GCS) plays central roles in C1 and amino acids metabolisms and the biosynthesis of purines and nucleotides. Manipulations of GCS are desired to promote plant growth or to treat serious pathophysiological processes such as aging, obesity and cancers. Reversed GCS reactions form the core of the reductive glycine pathway (rGP), one of the most promising pathway for the assimilation of formate and CO_2_ in the emerging C1-synthetic biology. H-protein, one of the four GCS component proteins (H, T, P and L) is generally considered a shuttle protein interacting with the other three proteins via a lipoyl swinging arm. Here, we discovered that without P-, T- and L-proteins, H-protein alone can catalyze GCS reactions in both glycine cleavage and synthesis directions *in vitro*. The surprising catalytic activities are related to a structural region of H-protein which can be manipulated. The results have impacts on engineering GCS to treat related diseases, to improve photorespiration, and to efficiently use C1-carbon for biosynthesis.

## Introduction

In the mitochondria of plant and animal cells as well as in the cytosol of many bacteria, the glycine cleavage system (GCS) comprising four (H, T, P and L) proteins catalyzes the reversible decarboxylation and deamination of glycine to yield CO_2_, NH_3_ and provide a methylene group for the conversion of tetrahydrofolate (THF) to N^5^,N^10^-methylene-tetrahydrofolate (5,10-CH_2_-THF)^1, 2, 3^. The overall reaction cycle catalyzed by GCS comprises three steps as illustrated in **Figure 1a** (hereinafter GCS is used to refer the four-enzyme system regardless of reaction direction). The reaction is first catalyzed by P-protein (glycine decarboxylase; EC 1.4.4.2) to yield CO_2_ from glycine and methylamine-loaded H-protein (H_int_) from the oxidized form (H_ox_) of the lipoylated H-protein (H_lip_). T-protein (aminomethyltransferase; EC 2.1.2.10) then catalyzes the release of NH_3_ and transfer the methylene group from H_int_ to THF to form 5,10-CH_2_-THF, leaving dihydrolipoyl H-protein (H_red_). Finally, L-protein (dihydrolipoyl dehydrogenase; EC 1.8.1.4) catalyzes the oxidation of H_red_ to regenerate H_ox_ in the presence of NAD^+^. H-protein as a shuttle protein interacts with the other three GCS-proteins via a lipoyl swinging arm and plays central role in the GCS.

**Figure. 1.**
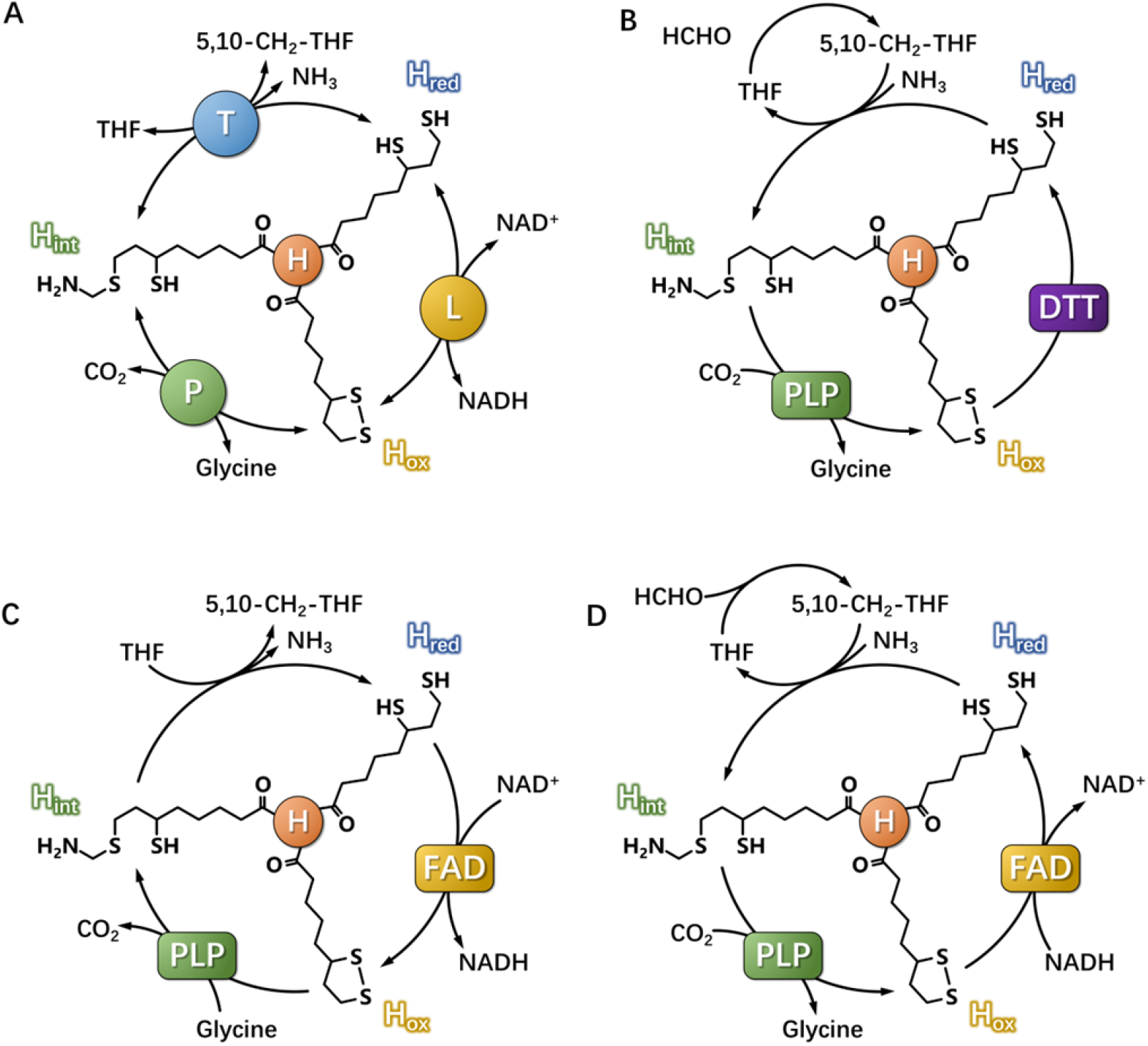
Schematic diagrams of the reversible glycine cleavage reaction catalyzed by GCS or stand-alone H-protein (H_lip_). (a) The glycine cleavage and synthesis reactions catalyzed by GCS with the complete set of enzymes; (b) Glycine synthesis reactions catalyzed by H_lip_ alone under the presence of PLP, THF and DTT (DTT as a reductant replacing the functions of FAD and NADH); (c) Glycine cleavage and (d) synthesis reactions catalyzed by H_lip_ alone under the presence of PLP, THF, FAD and NADH.

The physiological roles of GCS in various organisms have been well studied. In human and most vertebrates, GCS is part of the serine and glycine metabolism pathway. Serine is catalyzed by serine hydroxymethyltransferase (SHMT) to form glycine and 5,10-CH_2_-THF, and then the product glycine is degraded by GCS. The final product, 5,10-CH_2_-THF, whose methylene group derived from the β-carbon of serine or the α-carbon of glycine, is one of the few C1 donors in the biosynthesis process, such as the biosynthesis of purine and methionine^2^. Decrease or loss in the activity of GCS will lead to glycine accumulation in human body, which is linked to glycine encephalopathy^4^ (also known as nonketotic hyperglycinemia). Relevant studies have shown that most patients with glycine encephalopathy have a P-protein deficiency, and the rest are caused by T-protein or H-protein deficiency^5^. Moreover, recent studies have shown that glycine metabolism is associated with tumorigenesis, and P-protein as a key factor regulates glycolysis and methylglyoxal production in cancer cells^6, 7^. In C3 plants, GCS is the key enzymatic system that deals with a large amount of glycine in mitochondria during the photorespiration, and the activity of GCS directly determines the growth rate of plants. Knockout of GCS gene is lethal to plants, which is relevant to impaired one-carbon metabolism^8^, whereas overexpression of L-protein^9^ or H-protein^10^ have been shown to improve photorespiration rates for further increasing biomass yield.

Although GCS in most organisms runs mostly in the direction of glycine cleavage, it catalyzes glycine synthesis in a few anaerobic bacteria such as *Clostridium acidiurici*^11^, *Eubacterium acidaminophilum*^12^ and *Arthrobacter globiformis*^13, 14^. The reversibility of GCS was first discovered in mitochondrial extract of rat liver^15^, in *A. globiformis*^16^ and in cock liver mitochondria^17^, and most of these studies were already carried out in 1960-1980s. Today, attributing to the reversibility of GCS, GCS gains renewed attention of researchers, because reductive glycine pathway (rGP), with GCS as its key component pathway, is considered to be the most promising synthetic pathway for the assimilation of formate and CO_2_ to produce pyruvate^18^, a key precursor that enters the central metabolic pathway for cell growth and biosynthesis. Sánchez-Andrea *et al*.^*19*^ discovered that rGP functions in an anaerobic sulphate-reducing bacterium *Desulfovibrio desulfuricans*, and stated that it represents the seventh natural CO_2_ fixation pathway. Recently, rGP has been successfully introduced into *E*.*coli*^20, 21, 22, 23^ for autotrophic growth on formate and CO_2_. At the same time, part of this pathway was successfully transferred into *Saccharomyces cerevisiae*^24^, *Cupriavidus necator*^*25*^ and *Clostridium pasteurianum*^*26*^. However, the flux of rGP is quite low, which limits the growth of microorganism. It has been pointed out that the reaction catalyzed by GCS is the rate-limiting step in rGP^21^. Therefore, it is particularly important to understand the catalytic mechanism of GCS for increasing the flux of rGP. Substantial progress has been made in understanding the catalytic properties of GCS, and H-protein is so far considered to function merely as a shuttle protein of the cofactor lipoic acid. Lipoic acid is attached by an amide linkage to the conserved lysine residue of H-protein at the 64^th^ position, and the lipoylated H-protein (H_lip_) plays a pivotal role acting as a mobile substrate which undergoes a cycle of reductive methylamination, methylamine transfer and electron transfer in the enzymatic cycle of GCS^27^.

In this work, we discovered that H_lip_ alone can enable the GCS reaction cycle in both glycine cleavage (**Figure 1b**) and synthesis directions (**Figures 1c and 1d**) in the absence of P-, T- and L-proteins. The formation of glycine from C1 compounds in the presence of suitable cofactors was demonstrated by choosing HCHO as the source of α-carbon of glycine. More detailed analyses led to the striking finding that H_lip_ can apparently “catalyze” all the GCS reaction steps previously believed to be solely catalyzed by P, T and L-proteins, respectively. These findings not only shed new light into the functions of H-protein, but also provide useful hints for engineering H-protein and GCS, either for treating diseases such as hyperglycinemia, for enhancing biomass yield in plants, or for developing synthetic pathways for technical use of C1-carbons. The fact that stand-alone H_lip_ can catalyze the synthesis of the basic amino acid glycine from inorganic compounds may also have important implications for the evolution of life.

## Results

### Effects of components of the GCS reaction system on glycine cleavage and synthesis

On the basis of previous studies^8, 28^, we successfully constructed GCS catalyzed glycine cleavage and synthesis reactions *in vitro*. Normally, all the four GCS enzymes are included in the reaction system. During kinetic studies, we found that the reactions of glycine cleavage and glycine synthesis can also occur in the absence of certain GCS enzymes and reaction components. This triggered us to systematically examine the effects of missing a certain component or enzyme in the reaction mixture on the reaction rate of both reaction directions. As shown in **Table 1**, the lack of a certain component or enzyme can cause very different changes of the reaction rate. As expected, the reaction did not occur in the absence of essential substrates (glycine in the cleavage direction and NH_4_HCO_3_ in the synthesis direction). The presence of the H_ox_ was also vital, as no reaction was observed in the absence of H_ox_. However, varied reaction rates (10-76 % of the reference values) were observed when only one of the P-, T- and L-proteins was missing. Compared to the effects of GCS proteins, the missing of substrates and cofactors (THF, PLP, NAD or NADH) showed often stronger effects on the cleavage and synthesis of glycine. In this context, the effects of PLP were surprising: (1) missing of both P-protein and PLP resulted in neither cleavage nor synthesis of glycine; (2) while missing PLP alone resulted in strongly impaired glycine synthesis, it had, however, no negative effect on glycine cleavage. This might be partially explained by the fact that PLP is covalently bound to P-protein^29, 30, 31^. Therefore, P-protein expressed in *E. coli* might have PLP covalently bound to it during its expression. The purified P-protein might still have PLP attached to it and can therefore function well in decarboxylation without externally adding PLP. On the contrary, the effect of PLP absence was even worse than the absence of P-protein for glycine synthesis which implied the importance of PLP for the stand-alone catalytic activity of H_lip_.

**Table 1.** Effects of missing a certain component of the GCS reaction system on the rates of glycine cleavage (determined as HCHO formation from the degradation of 5,10-CH_2_-THF) and glycine synthesis.

| Missing component | Glycine cleavage reaction |  | Glycine synthesis reaction |  |
| --- | --- | --- | --- | --- |
| | ($\mu\text{M}$ HCHO $\cdot \text{min}^{-1}$) | GCS/ % | ($\mu\text{M}$ glycine $\cdot \text{min}^{-1}$) | rGCS/ % |
| None (Reference) | 22.48±3.47 | 100.00 | 5.95±0.13 | 100.00 |
| P-protein | 2.32±0.52 | 10.34 | 2.03±0.20 | 34.07 |
| T-protein | 11.67±0.42 | 51.91 | 4.55±0.16 | 76.53 |
| L-protein | 8.44±0.57 | 37.55 | 4.45±0.15 | 74.78 |
| H<sub>ox</sub> | 0.00 | 0.00 | 0.00 | 0.00 |
| P-protein+PLP | 0.00 | 0.00 | 0.00 | 0.00 |
| PLP | 24.92±2.67 | 110.86 | 0.98±0.13 | 16.42 |
| T-protein+THF | 1.11±0.33 | 4.93 | 0.43±0.08 | 7.20 |
| THF | 0.87±0.15 | 3.88 | 0.53±0.04 | 8.94 |
| NAD<sup>+</sup>/NADH | 4.88±1.54 | 21.73 | 5.76±0.05 | 96.72 |
| Glycine | 0.00 | 0.00 | - | - |
| NH<sub>4</sub>HCO<sub>3</sub> | - | - | 0.00 | 0.00 |
| HCHO | - | - | 0.98±0.02 | 16.42 |

### H_lip_ alone enables glycine synthesis and glycine cleavage reactions

The results in **Table 1** suggested that P-protein, T-protein and L-protein are not essential for the functionality of GCS both in glycine cleavage and glycine synthesis directions. This led us to the question if H_lip_ alone can “catalyze” glycine formation from NH_4_HCO_3_ and HCHO, or glycine cleavage in the opposite direction.

For glycine synthesis, the experimental results with H_lip_ as the only GCS protein in an array of reaction mixtures are presented in **Figure 2a**. Compared with the glycine synthesis catalyzed by the all four GCS proteins (GCS as control), the reaction rate catalyzed by H_lip_ alone at the same concentration of 10 µM (H-10) was somewhat lower, but glycine formation was well detected. With the increase of H_ox_ concentration the initial reaction rate increased and the final concentration of glycine synthesized was higher than that of the control. The above results prove that H_lip_ alone can apparently catalyze the synthesis of glycine from NH_4_HCO_3_ and HCHO in the presence of THF, PLP and DTT. To get the optimum reaction conditions for the glycine synthesis, the reaction rate of glycine synthesis catalyzed by H_lip_ alone was investigated under conditions of using different buffers, temperatures and pHs. **Figure 2b** shows the effect of different types of buffer, in which the order of the catalytic ability of H_lip_ was as follows: Tris-HCl≈Mops > HEPES > PBS. The effect of temperature and pH on the activity were studied by changing temperature from 4 °C to 95 °C (**Figure 2c**), and pH from 6.0 to 10.0 (**Figure 2d**). The reaction rate decreased sharply when the temperature was higher than 42 °C or pH was higher than 8.0. The optimum temperature and pH were at 37-42 °C and 7.5-8.0, respectively.

**Figure 2.**
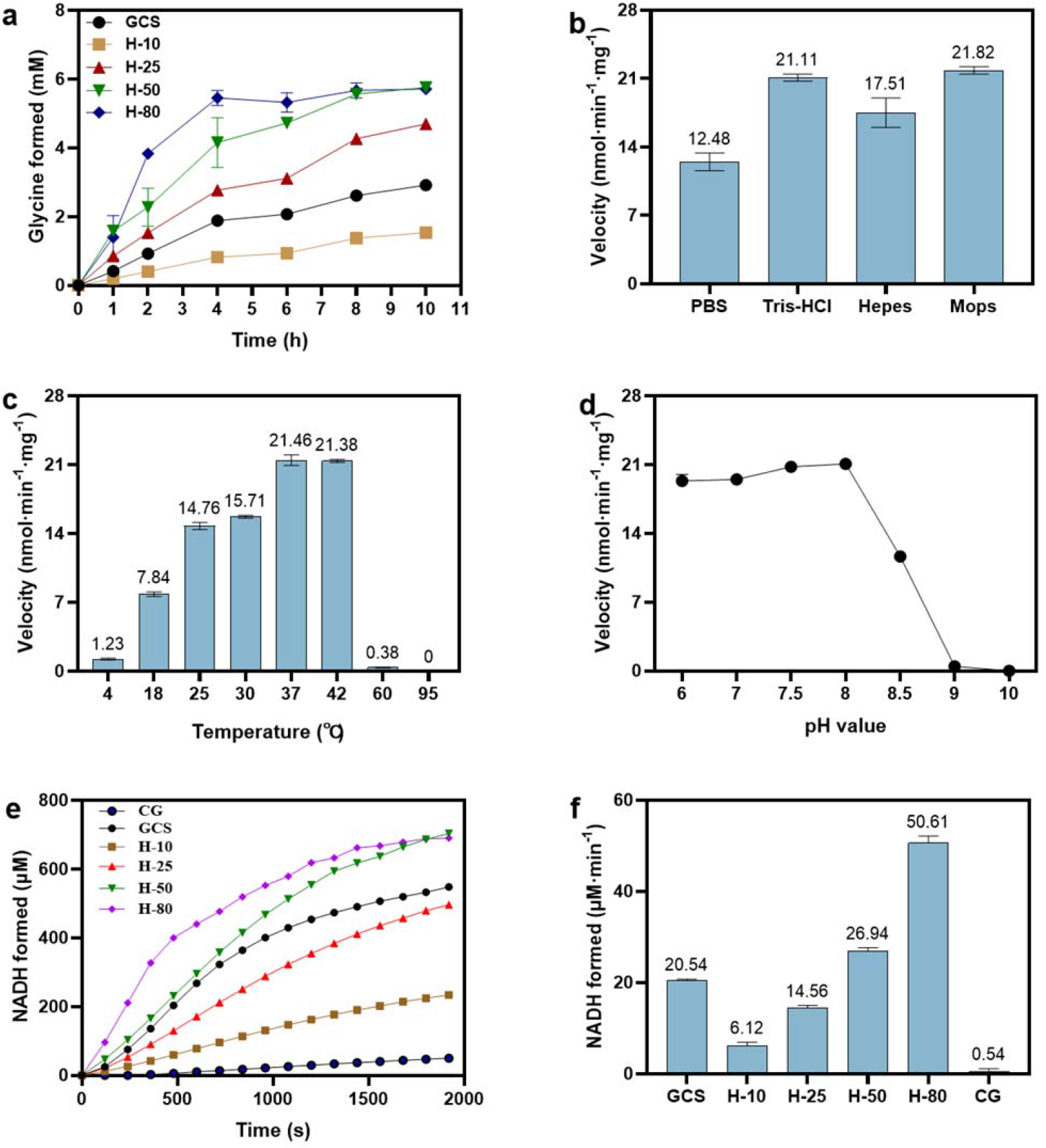
H_lip_ alone enabled glycine synthesis and glycine cleavage. Effects of H_ox_ concentration (a), buffer (b), temperature (c) and pH (d) on glycine synthesis. “GCS” refers to a reaction mixture for glycine synthesis as specified in “Materials and Methods” without missing any reaction components and enzymes; “H-10”, “H-25”, “H-50” and “H-80” were the same reaction mixture containing no P-, T- and L-proteins but only H_ox_ at 10 µM, 25 µM, 50 µM and 80 µM, respectively. (e) Effects of H_ox_ concentration on glycine cleavage. “CG” refers to no GCS enzymes in the reaction mixture, “GCS” refers to a reaction mixture for glycine cleavage as specified in “Materials and Methods” without missing any reaction components and enzymes; “H-10”, “H-25”, “H-50” and “H-80” were the same reaction mixture containing no P-, T- and L-proteins but only H_ox_ at 10 µM, 25 µM, 50 µM and 80 µM, respectively. (f) NADH formation rate in glycine cleavage catalyzed by different concentrations of H_ox_.

For glycine cleavage, the reaction could not be observed even at high H_lip_ concentrations (up to 80 µM) using the same reaction mixture as used for the GCS-catalyzed glycine cleavage reaction but without P-, T- and L-proteins. Later, we found out that when FAD, the coenzyme of L-protein, was added, H_lip_ alone was indeed able to activate the glycine cleavage, and the reaction rate increased with the increase of H_lip_ concentration, as shown by the time-courses of NADH formation (**Figure 2e**) and initial rates of glycine cleavage (**Figure 2f**). The essentiality of FAD for the glycine cleavage but not for glycine synthesis catalyzed by stand-alone H_lip_ is due to the presence of DTT which can convert H_ox_ to H_red_ required in the direction of glycine synthesis (details see below).

H-protein is a small heat-stable protein, so heating does not lead to precipitation. In literature, thermal stability of H-protein is therefore used to terminate the lipoylation of H-protein catalyzed by the enzyme lipoate-protein ligase A (LplA), in which LplA is completely denatured and precipitated ^32, 33^. We have tried to use heated H_ox_ (at 95 °C for 5 min) to catalyze the reactions of glycine synthesis and cleavage, but no reaction in either direction was observed (the details are discussed in a later section). We therefore speculate that the structure of H_lip_ was altered by heating at high temperature, which made it lose its catalytic activity shown above for glycine synthesis and cleavage.

In order to explore the reasons behind the function of the stand-alone H_lip_ observed, we further studied the effect of H_lip_ alone on the three GCS reaction steps, i.e., the glycine decarboxylation reaction (accompanied by the reductive aminomethylation of H_ox_ to H_int_) in the absence of P-protein, the aminomethyl transfer reaction in the absence of T-protein, and the electron transfer reaction without the presence of L-protein, respectively.

### Decarboxylation and carboxylation reactions in the absence of P-protein

The results in **Table 1** revealed that no activity could be measured for either the cleavage or the synthesis of glycine, when both P-protein and PLP were absent in the reaction mixtures. However, activities were observed when only P-protein was missing. We therefore speculate that the presence of PLP alone might be sufficient to enable H_lip_ to catalyze the decarboxylation/carboxylation reaction normally catalyzed by P-protein. This was confirmed by the HPLC results shown in **Figure 3a** for the decarboxylation reaction in the glycine cleavage direction. H_int_ was formed from H_ox_ without the presence of P-protein. This astonishing result suggests that glycine decarboxylation activated by H_lip_ alone can occur independent of P-protein, as long as PLP is present (**Figure 3a**) under the experimental conditions of this study.

**Figure 3.**
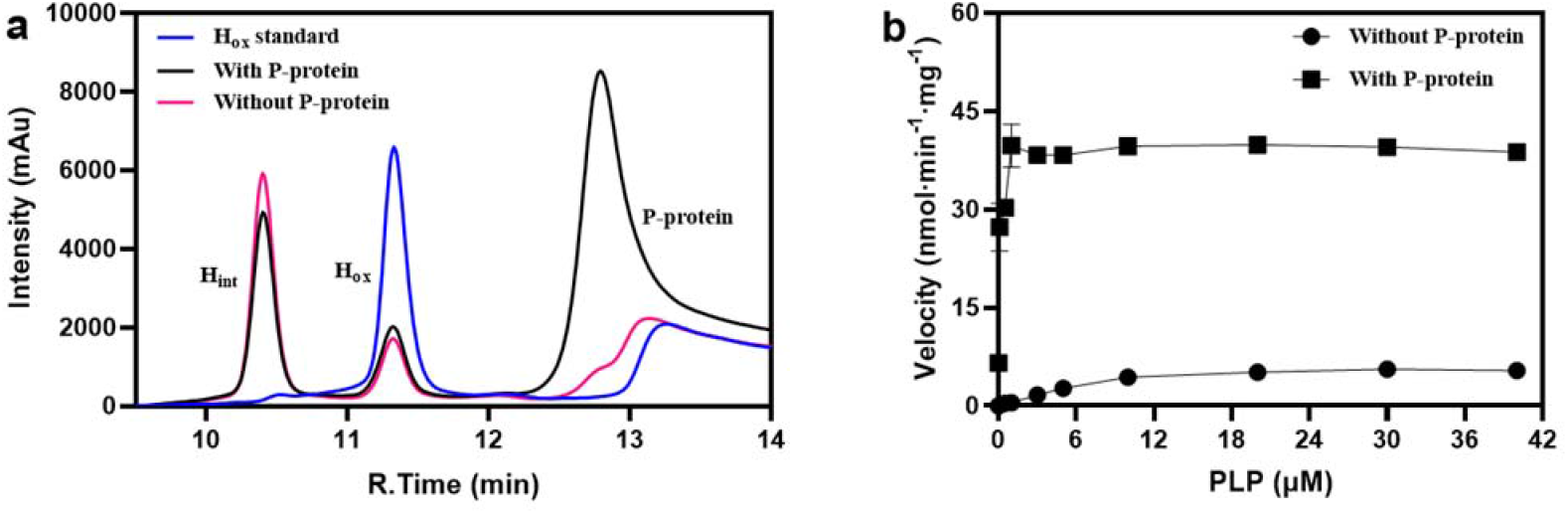
Examination of H_lip_ (H_ox_) for the function of P-protein. (**a**) Formation of H_int_ from H_ox_ during the glycine decarboxylation reaction was determined using HPLC to show that H_lip_ functions as decarboxylase in the presence of PLP. Chromatograph “With P-protein” refers to a reaction mixture containing 50 mM glycine, 50 µM H_ox_, 25 µM PLP, and 5 µM P-protein; Chromatograph “Without P-protein” refers to a reaction mixture similar to that “with P-protein” but without the addition of P-protein; Chromatograph “H_ox_ standard” refers to a test solution containing only H_ox_; (**b**) PLP-dependent glycine formation was determined to show that carboxylation can take place in the absence of P-protein, albeit at lower reaction rate. “With P-protein” refers to a reaction mixture for glycine synthesis as specified in “Materials and Methods” without missing any reaction components and enzymes, “Without P-protein” refers to a reaction mixture containing all reaction components and enzymes except for P-protein.

For the carboxylation reaction in the glycine synthesis direction, in the absence of P-protein, glycine formation could be still detected and the reaction rate showed a nearly linear increase with the PLP concentration in the low PLP concentration range, albeit that the reaction rate was lower than those determined in the presence of P-protein (**Figure 3b**). This is understood that CO_2_ fixation (carboxylation) has a higher energy barrier and P-protein is needed for its activation.

### Aminomethyl transfer reaction in the absence of T-protein

According to the results in **Table 1**, the overall GCS reaction in both glycine cleavage and glycine synthesis directions could still precede reasonably well in the absence of T-protein in the reaction mixtures. In comparison, it is obvious that the absence of THF had a more significant negative effect on the reaction rate regardless of the reaction directions, i.e. with a reduction of the reaction rate for over 96 % in glycine cleavage and 91 % in glycine synthesis. **Figure 4a** shows the change of NADH production with time in the direction of glycine cleavage under different conditions. The initial rate without adding T-protein was still more than half of that with adding T-protein, whereas in the experimental group “without THF” the formation of NADH could be hardly detected. For the aminomethyl transfer reaction in the direction of glycine synthesis, **Figure 4b** shows that H_int_ could still be generated without adding T-protein in the reaction mixture. It was found by Kochi *et al*^14^ that when dihydrolipoic acid, HCHO and NH_4_^+^ were mixed together, compounds in the form of -S-CH_2_NH_2_ could be obtained. The question then arises: is the aminomethylation of H_red_ to H_int_ in the absence of T-protein the result of a complete non-enzymatic reaction due to the presence of HCHO and NH_4_Cl in which H-protein acts only as the shuttle protein?

**Figure 4.**
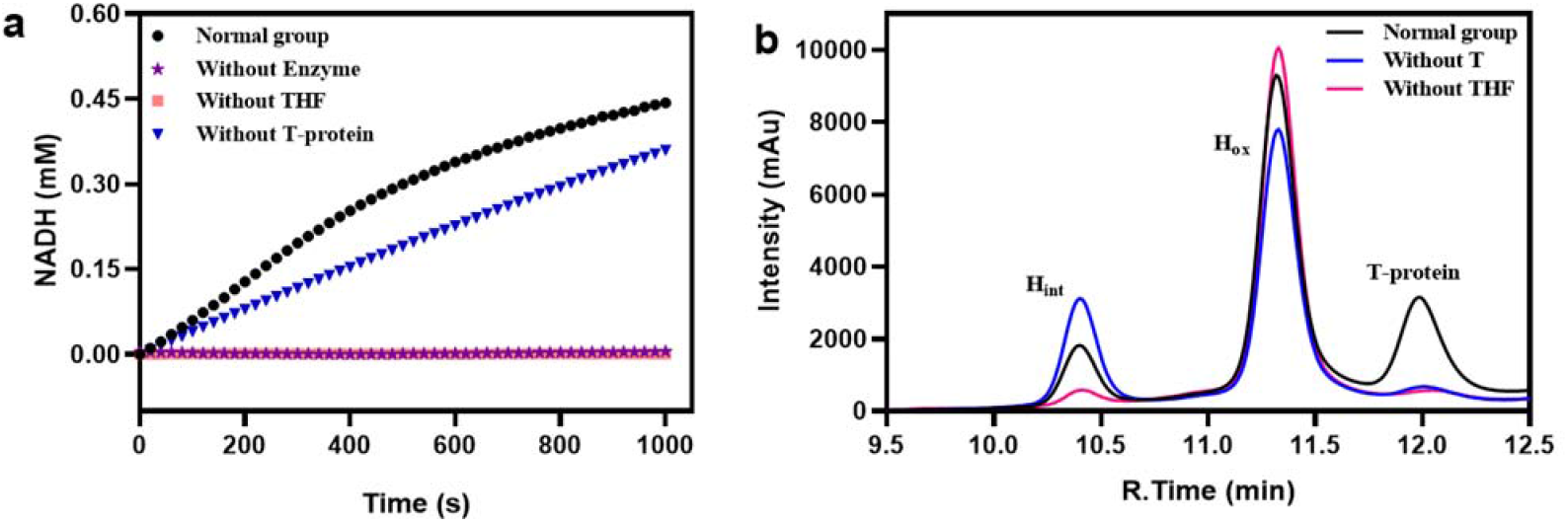
Examination of H_lip_ for the function of T-protein. (**a**) Effects of THF or T-protein absence on the overall glycine cleavage reaction rate. “Normal group” refers to a reaction mixture containing all reaction components and enzymes required; “Without Enzyme” refers to a reaction mixture containing all reaction components but no GCS enzymes; “Without THF” refers to a reaction mixture containing all reaction components and enzymes except for THF; “Without T-protein” refers to a reaction mixture containing all reaction components and enzymes except for T-protein. (**b**) Detection of H_int_ by HPLC proved that the aminomethyl transfer reaction can take place spontaneously. “Normal group” refers to a reaction mixture containing 50 µM H_ox_, 0.5 mM THF, 20 mM DTT, 50 mM NH4Cl, 10 mM HCHO, and 5 µM T-protein; The reaction mixtures of “Without T” and “Without THF” were the same as that of “Normal group”, but no T-protein or THF was added, respectively.

### Interconversion of H_ox_ and H_red_ in the absence of L-protein

The function of L-protein is to catalyze the interconversion between the H_ox_ and its reduced form dihydrolipoyl H-protein (H_red_) involving NAD^+^/NADH and FAD/FADH_2_. The experimental results in **Table 1** show that in the absence of L-protein both the glycine cleavage and synthesis systems can still work. Previous studies^34, 35^ have shown that the disulfide bond-reducing agent TCEP allows the reduction of the disulfide bond of the lipoyl group associated with the H-protein during the course of the reaction catalyzed by L-protein. Since a certain amount of DTT was added to prevent the oxidation of THF during the preparation of THF stock solution, we expected that DTT should have the same reduction function as TCEP. **Figure 5a** schematically shows the interconversion between H_ox_ and H_red_ through the combination of the reduction of H_ox_ by TCEP (Left) or DTT (Right) with the re-oxidation of H_red_ catalyzed by L-protein, yielding thereby NADH from added NAD^+^. The functionality of this combination was demonstrated by the relevant results presented in **Figure 5b**. Next, we compared the effects of three different reducing agents (TCEP, DTT and β-ME) on the reducibility of H_ox_ to H_red_ in different buffers (Tris-HCl, PBS, Mops and HEPES) (**Figure 5c**). It was found that the reducibility of H_ox_ by DTT in four different buffers is PBS > Mops > Tris-HCl > HEPES; TCEP had no reducing effect on H_ox_ in Tris-HCl but showed the strongest reducing power in PBS, even much better than DTT. β-ME was obviously not suitable to reduce H_ox_ in these four buffers. Thus, for glycine synthesis, the functionality of L-protein can be well replaced by DTT or TCEP, as also evidenced by the result shown in **Table 1** that the absence of NAD^+^/NADH did not affect glycine synthesis.

**Figure 5.**
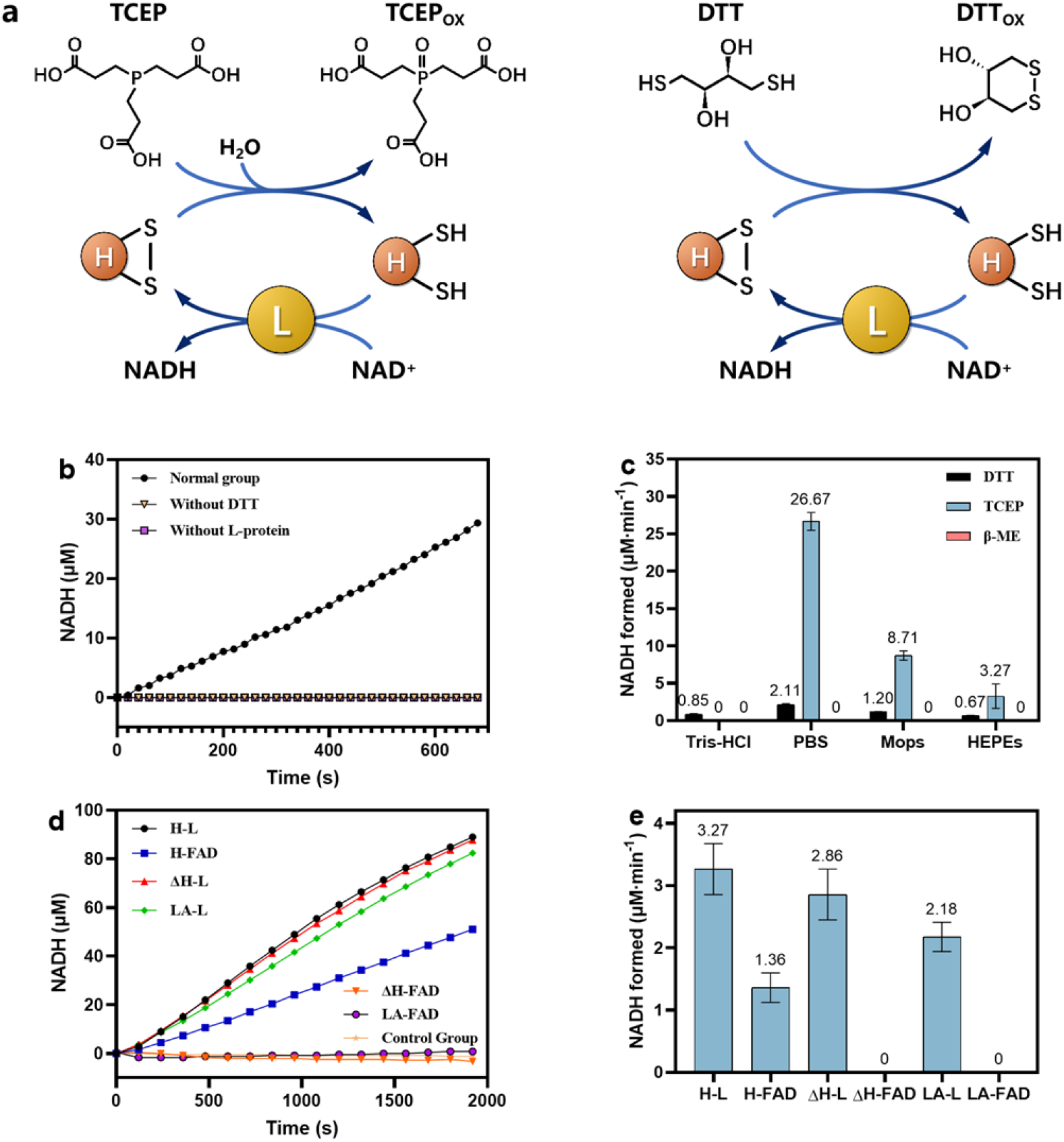
Examination of H_lip_ for the function of L-protein. (**a**) Interconversion between H_ox_ and H_red_ by combining the reduction of H_ox_ to H_red_ by TCEP (Left) or DTT (Right) with the re-oxidation of H_red_ to H_ox_ catalyzed by L-protein. (**b**) Reduction of the lipoamide group of H_ox_ by DTT. “Normal group” refers to a reaction mixture for electron transfer reaction as specified in “Materials and Methods” without missing any reaction components and enzymes. “Without DTT” refers to a reaction mixture similar to “Normal group” but without adding DTT. “Without L-protein” refers to a reaction mixture similar to “Normal group” but without adding L-protein; (**c**) Comparison of different disulfide reductants on the reduction of H_ox_ in different buffer solutions. (**d**) Time courses of NADH formation as the result of the redox reaction of H_lip_ (H) or heated H_olip_ (ΔH) or lipoic acid (LA) in the presence of either L-protein (L) or FAD. (**e**) NADH formation rates in the redox reaction of H_ox_, heated H_ox_ and lipoic acid in the presence of either L-protein or FAD.

In order to verify that H_lip_ has the catalytic function of L-protein, we used FAD, the redox coenzyme of L-protein, instead of L-protein in combination with DTT to observe whether the redox reaction can still occur. To this end, NADH formation as the result of the redox reaction of H_lip_ (**Figure 5a**) by either L-protein or FAD was measured. Considering the results shown in **Figure 5d** and **Figure 5e**, the following conclusions can be drawn: (1) As long as L-protein is present, the redox reaction of the disulfide bond on the lipoyl group is comparable, with the lipoyl group bound to H-protein (H_lip_) showing a slightly higher activity than that not bound (in lipoic acid). (2) In the absence of L-protein, the redox reaction of the lipoyl group bound to H-protein still occurred with the help of FAD, though to a lesser extent, but FAD was not able to replace L-protein in the reoxidation of dihydrolipoic acid, indicating that binding of the lipoyl group on H-protein is the prerequisite for the function of FAD in the absence of L-protein. To find out whether this is simply due to “fixation effect” of lipoyl group bound to H-protein which enables an easier approach of FAD to the lipoyl group, or this is facilitated through an unknown interaction of FAD with H-protein, we also examined to use heat-treated (95 °C for 5 min) H_ox_, which still had the lipoyl arm linked to it. Interestingly, in the presence of L-protein, there was nearly no difference in the redox reaction of the lipoyl group between the heated H_ox_ and the unheated H_ox_, however, FAD completely lost its function on the heated H_ox_, clearly suggesting that an interaction of FAD with H-protein is required for its function in the absence of L-protein, and heating-induced structural change of H-protein destroyed the possibility of such interaction.

### Possible reasons of apparent catalytic functions of H_lip_ in glycine cleavage and glycine synthesis

The above results show that for *in vitro* GCS reactions, H_lip_ alone enables both the glycine synthesis and the glycine cleavage without the presence of P-, T-, and L-proteins. It seems that H_lip_ might functionally replace at least P- and L-proteins and acts as glycine carboxylase and dihydrolipoyl dehydrogenase with the help of PLP and FAD, respectively. It is also suggested by the experiment results shown in **Figure 5d** and **5e** that heated H_ox_ lost the catalytic function of L-protein, obviously because heated H_ox_ cannot interact with FAD for the redox reaction of the lipoyl group bound to H-protein.

We therefore further systematically studied the effects of heating (95 °C for 5 min) on the catalytic activity of H_lip_ for glycine synthesis, in comparison to unheated H_ox_ as well as to lipoic acid. By either using H_lip_ alone or combined with other GCS enzymes, the overall reaction rate of glycine synthesis was measured. From the results in **Table 2**, we can ascertain several interesting observations and conclusions. First, the overall glycine synthesis reaction could be catalyzed by the unheated H_ox_ alone; adding P-protein significantly enhanced glycine synthesis, showing the importance of P-protein; the addition of either T- or L-protein has no positive effect. In fact, the *in vitro* glycine synthesis could run even better without T- and L-proteins, suggesting that in the presence of H_lip_ the two individual steps, namely aminomethyl transfer and electron transfer catalyzed by the two proteins respectively, could take place through spontaneous aminomethylation of H_red_ to H_int_ in the presence of HCHO and NH_4_^+^ and reduction of H_lip_ to H_red_ by DTT. Second, compared with the unheated H_ox_, the heated H_ox_ alone could not catalyze the reaction of glycine synthesis; adding P-protein and L-protein partially revived glycine synthesis to different extents, indicating that heated H-protein lost the catalytic ability but was still functional as the shuttle protein of lipoyl group; the addition of T-protein did not bring any effect. Furthermore, to our surprise, when the heated H_ox_ was added with the other three GCS proteins to form a complete GCS, the rate of glycine synthesis was even higher than the GCS containing the unheated H_ox_, indicating that heated H_ox_ loses its catalytic function but the heat-induced change is even beneficial for H-protein to exert its role as a lipoyl-carrying protein to work together with the other three GCS proteins. Finally, in the presence of P-, T- and L-proteins, free lipoic acid can act as intermediate substrate to sustain the reversed GCS reaction towards gylcine synthesis under the given experimental conditions, even though to a much lower extent than that observed in the case of lipoyl group bound to H-protein.

**Table 2.**
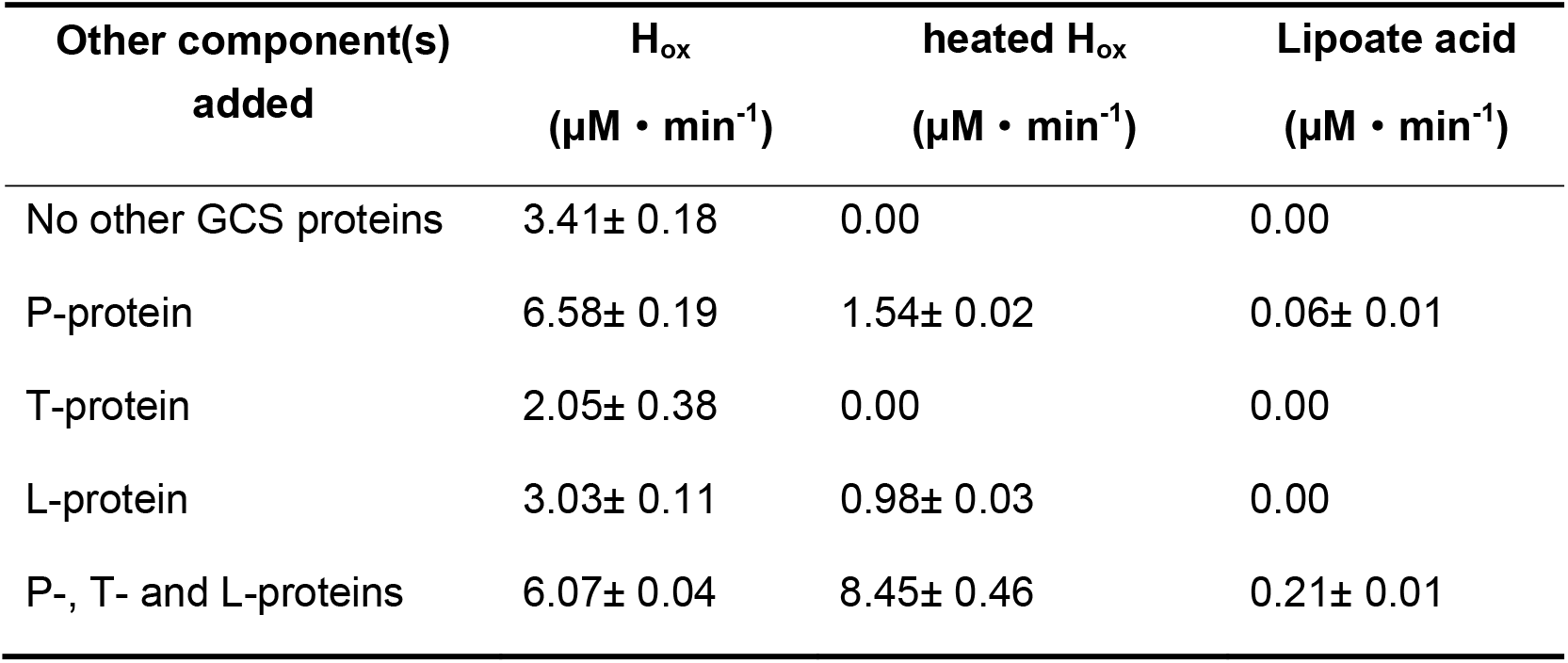
Synthesis of glycine using unheated H_ox_ or heated H_ox_ or lipoic acid at the same concentration of 10 µM, either alone or in varied combination with other GCS enzymes

Heating H_lip_ led to the loss of its catalytic activity regarding glycine synthesis. Although we found no change of the HPLC retention time of H_lip_ in the reversed-phase HPLC chromatograph after heating, which indicates no obvious change in the overall polarity and size of H_lip_, heating may induce structural changes that are vital for the catalytic activity of H_lip_. We therefore additionally performed HPLC analysis of H_int_ and H_ox_ to further determine the catalytic activity of H_lip_ in the two individual reaction steps normally catalyzed by P-protein and T-protein, respectively. As shown in **Figure 6a**, in the group of unheated H_ox_ the formation of H_int_ from H_ox_ clearly demonstrated that H_lip_ was not only a lipoyl-carring protein but could also replace P-protein in catalyzing the glycine decarboxylation reaction. H_int_ was not detected in the group of heated H_ox_, indicating that heated H_ox_ lost the catalytic activity of P-protein. **Figure 6b** shows the reaction results of aminomethyl transfer from H_red_ (generated *in situ* from H_lip_) to H_int_. The group of unheated H_ox_ clearly exhibited the catalytic activity of T-protein; in the group of heated H_ox_ there was a small amount of H_int_ formation found, however, this low level of aminomethyl transfer activity was most likely not due to a catalytic activity of T-protein manifested by heated H_ox_, but due to a spontaneously occurring aminomethyl transfer reaction. Since heated H_ox_ still preserves its function as shuttle protein, the difference between the unheated and heated H_ox_ in the aminomethyl transfer reaction may provide an answer to the question in section 3.4 that in the absence of T-protein, the participation of H_lip_ in the aminomethyl transfer reaction is not only as shuttle protein.

**Figure 6.**
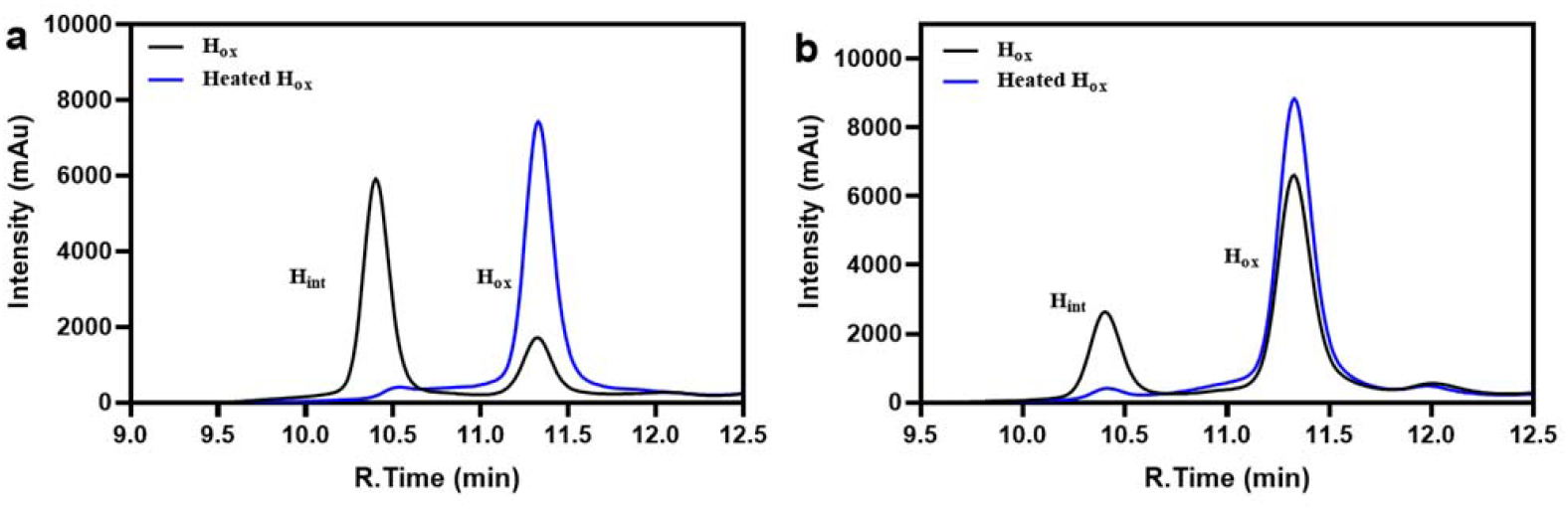
Effects of heating on the catalytic activity of H_lip_ as decarboxylase or aminomethyltransferase. (**a**) Decarboxylation reaction of glycine. “H_ox_” refers to a reaction mixture containing 50 mM glycine, 50 µM H_ox_, and 25 µM PLP; “heated H_ox_” was the same as “H_ox_” except for using heated H_ox_. (**b**) Aminomethyl transfer reaction (started from H_ox_, which was reduced to H_red_ by DTT, before H_red_ is converted to H_int_ by aminomethyl transfer reaction). “H_ox_” refers to a reaction mixture containing 50 µM H_ox_, 0.5 mM THF, 20 mM DTT, 50 mM NH_4_Cl, and 10 mM HCHO; “heated H_ox_” was the same as “H_ox_” except for using heated H-protein.

To explain the above experimental observations we examined the possibility that the catalytic ability of unheated H_ox_ is due to intermolecular interactions of H_lip_ itself, and such interactions would enable H_lip_ to catalyze the interconversion of the different forms (H_lip_, H_red_ and H_int_) to complete cycle by cycle the electron transfer, aminomethyl transfer, and reductive aminomethylation. In order to verify this hypothesis, we developed a concept to distinguish a possible intermolecular interaction of H_lip_ from its function as shuttle protein. On the one hand, the lysine residue at position 64 of H-protein for binding lipoic acid was mutated into alanine to obtain H_K64A_ mutant protein, which is unable to be lipoylated and consequently unable to act as shuttle protein, but is still expected to be able to uphold the capability of intermolecular interactions of the wild-type H-protein. On the other hand, as discussed above, heated H_ox_ losing its catalytic activity for glycine synthesis well preserves its function as a shuttle protein. Therefore, if the hypothesis holds true, we would expect that mixing heated H_ox_ with H_K64A_ would “restore” the catalytic activity exhibited by unheated H_ox_ towards glycine synthesis. No glycine formation could be detected which rules out the hypothesis and points out that the interconversion of the three forms of lipoylated H-protein, and consequently the stand-alone catalytic activity of H-protein, is not the result of intermolecular interactions of H-protein itself.

An answer to the question of why heating destroys the catalytic ability of H-protein but still preserves its function as shuttle protein might lie in the special surface structure of H-protein. As shown by the crystal structure of H_int_ (**Figure 7a**), following methylamine transfer the lipoamide-methylamine arm enters into the hydrophobic cavity on the surface of the H-protein and prevents thereby it from nucleophilic attack by water molecules^28, 36, 37, 38^. Based on our previously molecular dynamic simulation study, the lipoamide-methylamine arm may interact with some amino acid residues in the proximity of the cavity^39^, such as Glu-12, Glu-14, Ser-67, Cys-68 and Tyr-70 (**Figure 7b**). We speculated that the catalytic activity found for the stand-alone H_lip_ is related to the structure of the cavity. Therefore, to verify this assumption, the above five residues of the wild-type H-protein were mutated to alanine. Assays of glycine synthesis were then performed with these mutated H_lip_ in comparison with the wild-type H_lip_. As shown in **Figure 7c**, compared to the wild-type H_lip_ the glycine synthesis rates were strongly reduced in reaction mixtures containing H_lip_ mutants. However, as shown in **Figure 7d**, when these mutants were combined with the other three GCS proteins, the glycine synthesis rates of all the mutants were increased to levels comparable or even better than that of the wild-type H_lip_. The results are very similar to what observed with heated H_ox_ (**Table 2**). It is therefore confirmed that the cavity on the H-protein surface plays a decisive role in the catalytic functions of H-protein, and alterations of the cavity structure (in size or form) either through mutation or heating will reduce or even destroy the stand-alone catalytic functions of H-protein (**Figure 7c, Table 2**) due to yet unclear mechanisms. One possible reason might be that the lipoamide-methylamine arm cannot properly enter the deformed cavity, resulting in the failure or imbalance of the GCS cycle in the presence of heated H_ox_ alone. By adding other GCS proteins, H-protein is mainly required to act as a shuttle protein and, consequently, GCS reactions can be revived (**Table 2**). Moreover, under the given *in vitro* reaction conditions, the lipoamide-methylamine arm may undergo fast and continuous reaction in the GCS cycle that not only minimizes the probability of its hydrolysis, but also maintains or even increases the reaction efficiency by omitting the process of its entry and exit from the cavity (**Figure 7d, Table 2**).

**Figure 7.**
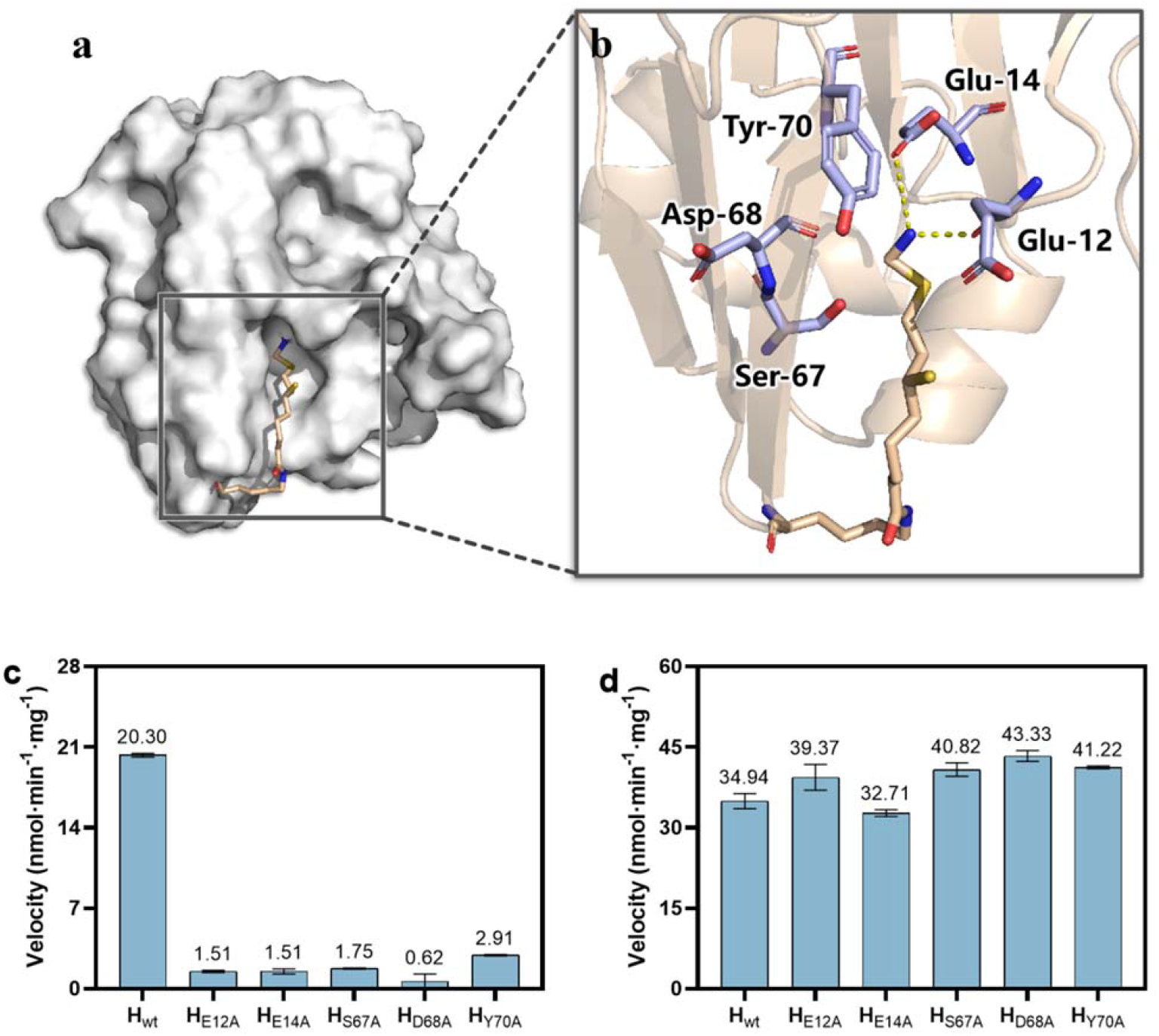
Study of the essential roles of the H-protein cavity. (**a**) Three-dimensional structure of *E. coli* H-protein bearing the lipoamide-methylamine arm protected in the cavity (ecH_int_). (**b**) Docking of the lipoamide-methylamine arm in the cavity with highlighting of some surrounding amino acid residues selected for mutation. (**c**) The rate of glycine synthesis catalyzed by stand-alone H_lip_ or its mutants. (d) The rate of glycine synthesis catalyzed by GCS comprising H_lip_ or its mutants and the other three GCS proteins.

## Discussion

In this work, we show for the first time that stand-alone lipoylated H-protein (H_lip_) has the catalytic functions so far believed to be carried out by the P-, T- and L-proteins of GCS. It enables glycine cleavage reactions, as well as the reversed reactions towards glycine synthesis with NH_4_HCO_3_ and HCHO as the substrates. The K_cat_ value for the overall synthesis reaction is about 0.01 s^−1^ for GCS catalyzed reaction and 0.0057 s^−1^ for H-protein alone catalyzed reaction. After purification of the GCS proteins, we used the most commonly used methods SDS-PAGE and HPLC to verify that there was no obvious residual of other GCS proteins in the purified H-protein solutions, though we did not confirm this by using more precise methods like mass spectroscopy. Through calculations, we can state that even if other GCS proteins would exist in the H-protein solution (e.g. up to 10%), it will not qualitatively affect the main conclusions drawn in our work (see Supplementary Materials for detailed explanation). Therefore, the purity of H-protein meets the requirement needed for this study.

The stand-alone catalytic activity of H_lip_ is closely related to the cavity on the H-protein surface, where the lipoyl swing arm is bound. Both heating H_lip_ and mutating cavity-related amino acid residues result in complete loss or strong reduction of the stand-alone catalytic activity of H_lip_, because they may cause deformation of the cavity, resulting in failure of the lipoamide-methylamine arm to properly enter the cavity and consequently failure of GCS reactions. Cohen-Addad *et al*.^*36*^ suggested that the lipoamide-methylamine arm is locked into a very stable configuration within the hydrophobic cavity and therefore highly stable against the non-enzymatic hydrolysis (which leads to the release NH_3_ and HCHO) due to nucleophilic attack by water molecules.

However, our experiments surprisingly show that when heated H_ox_ or a H_ox_ mutant was combined with the other GCS proteins, the rate of glycine synthesis was recovered or even increased (**Table 2** and **Figure 7**). This implies that under the given *in vitro* reaction conditions, heated H_ox_ truly acts only as shuttle protein in the presence of P-, T- and L-proteins, it is unnecessary and even disadvantageous for the lipoamide-methylamine arm to enter the cavity in order to undergo GCS reactions. Indeed, there was no obvious decrease of the peak area of H_int_ on HPLC even after hours of waiting, indicating that the hydrolysis of H_int_ is very slow (data not shown). This is in consistence with our recent finding that mutations of the key residue Ser-67 which reduce the residence time of the lipoamide-methylamine arm in the cavity can significantly increase the *in vitro* GCS activity^39^. The molecular dynamic simulations of H-protein carried out by our group^39^ also implies that the lipoamide-methylamine arm can leave the cavity of unheated H-protein even without the interaction with T-protein. This is confirmed by further molecular dynamic simulations of H-protein (results not published yet). These results therefore raise the question why the lipoamide-methylamine arm should enter the cavity and be protected in the GCS system? Based on the fact that the cavity is closely related to the stand-alone catalytic activity of H-protein, perhaps we may make a bold speculation for the interpretation of these results from a perspective of evolution. *In vivo*, GCS may have evolved from a simple system for glycine cleavage catalyzed by H-protein alone at the early time of evolution to a sophisticated system, in which H-protein is assisted by specialized P-, T-, L-proteins for more effective catalysis of the GCS reactions to meet the growing metabolic requirements of organisms. However, the cavity structure and the stand-alone catalytic functions of H-protein have been retained till now.

Oliver^40^ discovered that the H-protein and the small subunit of ribulose-1,5-bisphosphat-carboxylase/-oxygenase (RuBisCo) have obvious similarities in plants. The two proteins are not only about the same size, but also have similar mechanism in terms of transcriptional control of the corresponding genes. It has been reported that there are striking sequence and structure similarity between H-protein and the E2 protein of pyruvate dehydrogenase complex (PDC)^27^. Therefore, further research on the catalytic mechanism of H-protein may give useful hints for understanding the evolution and function of PDC and other 2-oxoacid dehydrogenase multi-enzyme complexes (e.g. alpha-ketoglutarate dehydrogenase complex) which are all of fundamental importance in cellular metabolism, governing the synthesis of C1-C4 metabolites for life.

H_lip_ alone catalyzes the glycine cleavage and synthesis *in vitro* with the help of the cofactors PLP, THF and FAD. This seems in contradictory to the results reported in the previous literature that deletion in gcvP or gcvT is lethal for organisms^8, 41^. By comparing the differences between the in *vivo* and *in vitro* conditions, it is conceivable that *in vivo* these cofactors are stoichiometrically linked to P-, T and L-proteins, respectively, to play their dedicated catalytic roles in a concerted action with H-protein. The *in vivo* concentrations of these cofactors are much lower than what we used in our *in vitro* experiments to facilitate the cleavage or synthesis of glycine catalyzed by H-protein alone. Thus, at one hand, it may not be feasible for cells to maintain high concentration pools of these cofactors; at the other hand, it might be beneficial for cells to have the sophisticated complete GCS system for fast and fine tuning and adapting of this important system to the changes in metabolic demands. This may be one reason why P-, T-, L-proteins are necessary *in vivo*. This study shows that for the stand-alone catalytic functions of H_lip_ an interaction of FAD with H_lip_ is required for its function in the absence of L-protein, and heating-induced structural change of H_lip_ destroyed the possibility of such interaction (**Figure 5d**). It is most likely that PLP also needs to interact with H_lip_ to exert its function in the absence of P-protein. This thus raises questions: How do PLP and FAD interact with H_lip_? Is it similar to the binding of PLP to P-protein and FAD to L-protein? To answer these questions, more experiments will be carried out in the future studies.

In addition, the overall reaction rate of GCS with T-protein deficiency was only reduced to 52 % in the direction of glycine cleavage and 76.5 % for glycine synthesis, compared to those of the complete GCS, indicating that T-protein had the least effect on the catalytic activities of GCS. These results are in agreement with the results of Timm *et al*.^8^ who showed that knockdown mutants of Arabidopsis containing very low T-protein expression under physiological conditions were able to grow and propagate in normal air, only showing some minor changes. Meanwhile, their study also found that the knockout mutation without T-protein expression was lethal even in non-photorespiratory environment of air enriched to 1 % CO_2_. This result may indicate that THF needs weak binding with T-protein to participate in glycine cleavage and synthesis reaction *in vivo*. When the T-protein is completely knocked out, THF cannot participate in the reaction, resulting in slow reaction rate and plant death. The deficiency of THF is far more detrimental and has the greatest impact on the reaction rate of GCS *in vitro* (**Table 1**).

Although we have identified the cavity of the H-protein as the key structural region that determines the catalytic activities of stand-alone H_lip_, the specific catalytic mechanism is not explored from the perspective of structure and molecular interaction. By studying the structures and their dynamics of heated and unheated H-proteins, it will certainly help to better understand the mechanism. This in turn will help to better engineer H-protein and GCS, leading to new possibilities to improve the growth and physiology of cells, organisms and plants and to design industrial microbes for utilizing C1 compounds for biosynthesis.

## Materials and Methods

### Materials

The substrates glycine, nicotinamide adenine dinucleotide (NAD^+^, NADH), Tris (2-carboxyethyl) phosphine (TCEP) and the derivatization reagents dansyl chloride were purchased from Yuanye Bio-Technology (Shanghai, China). Dithiothreitol (DTT), β-mercaptoethanol (β-ME), pyridoxal 5’-phosphate monohydrate (PLP) and flavin adenine dinuleotide (FAD) were obtained from Aladdin (Shanghai, China). 6-(RS)-Tetrahydrofolate (THF) was obtained from Sigma-Aldrich (St. Louis, MO, USA). Other chemicals in this study were of analytical grade and purchased from Solarbio (Beijing, China) or Sinopharm (Shanghai, China), unless otherwise noted. *Escherichia coli* Top10 and BL21 (DE3) were used for plasmid construction and the overexpression of recombinant proteins, respectively. Ni^2+^-NTA resin was purchased from Genscript (Nanjing, China). Amicon^®^ Ultra-15 filtration devices (molecular size cut-off 10 KDa for H-protein, 30 KDa for T-protein and L-protein and 100 KDa for P-protein) were purchased from Millipore (Billerica, MA, USA). Mut Express II Fast Mutagenesis kit V2 was purchased from Vazyme (Jiangsu, China). BCA protein assay kit and sodium dodecyl sulfate-polyacrylamide gel electrophoresis (SDS-PAGE) gels were purchased from SolarBio (Beijing, China). Luria-Bertani (LB) medium containing tryptone (10 g/L), yeast extract (5 g/L) and NaCl (10 g/L) were used for cloning and expression, and tryptone and yeast extract were purchased from Oxoid.

### Enzyme preparation

The plasmids and bacterial strains used in this experiment were given in **Table 3**. Oligonucleotide sequences of primers used for cloning target proteins were given in Table 1 of Supplementary Materials. The genes coding for P-protein, T-protein, L-protein and H-protein were amplified from *E. coli* K12 genomic DNA, then cloned into the expression vector pET28a (NdeI and XhoI). *E*.*coli* BL21 (DE3) harboring the resulting constructs (pET28a-P, pET28a-T, pET28a-L and pET28a-H) were cultured in LB medium supplemented with 50 mg/L of kanamycin at 37°C until the OD_600_ of the culture reached 0.6-0.8, Isopropyl-beta-D-thiogalactopyranoside (IPTG) was added to a final concentration of 0.2 mM to induce protein expression for 12 h at 30 °C.

**Table 3.**
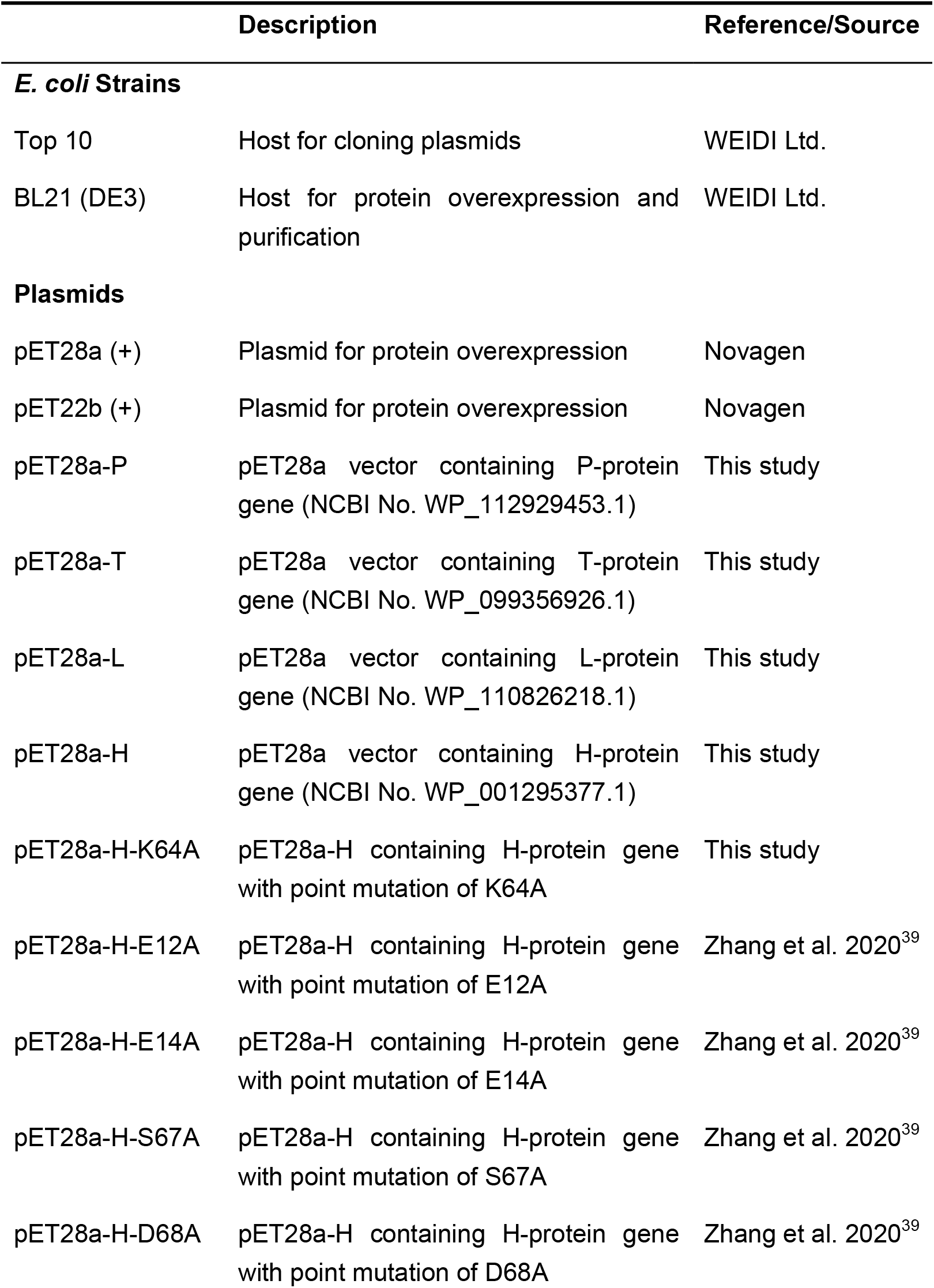
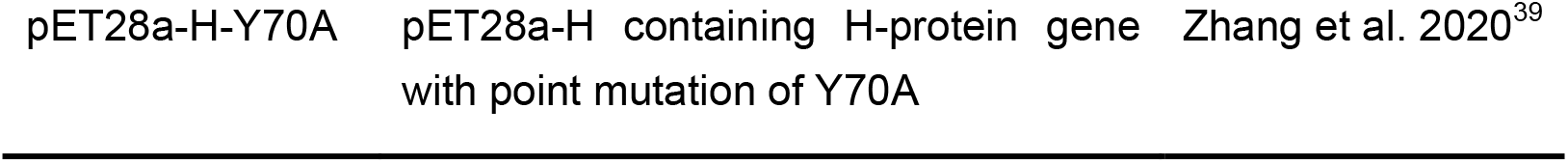
Strains and plasmids used in this study.

The plasmid pET28a-H was used as a template to generate mutations using Mut Express II Fast Mutagenesis kit V2^39^. Lipoylation of H-protein was performed during its over-expression *in vivo*. To this end, the strain containing the plasmid pET28a-H or a H-protein mutant were added with lipoic acid (200 μM, pH 7.0), prior to starting the cultivation to directly obtain lipoylated H-protein (H_lip_). Following the overexpression, enzymes were purified as described previously^42^. The purified enzymes were checked by SDS-PAGE (shown as Figure S1 in Supplementary Materials). In the lane of the H protein no residuals of T, P and L proteins were found except for the existence of low amount of the inactive H apo-protein (H_apo_). In addition, HPLC analysis also confirmed that there was no obvious residual of other GCS proteins in the purified H-protein solutions. H_lip_ obtained in such a way is considered to primarily exist in the oxidized form of H-protein (H_ox_). Whereas H_lip_ is mostly used to refer to lipoylated H-protein in general, it is numerically equivalent to H_ox_ in this work when concrete reactions involving H_lip_ are referred to and vice versa.

### Enzyme activity assays

#### The overall reaction of glycine cleavage

##### (1) Glycine cleavage catalyzed by GCS

The reaction mixture contained Tris-HCl (50 mM, pH 7.5), 0.5 mM THF, 20 mM DTT, 25 μM PLP, 5 mM NAD^+^, 5 µM P-protein, 5 µM T-protein, 5 µM L-protein and 10 µM H_ox_. After premixing and centrifugation, the reactions were initiated by the addition of 50 mM glycine, and carried out for 30 min at 37 °C. The overall GCS activity was determined by measuring either NADH formation at 340 nm using an Enspire multimode plate reader (PerkinElmer, USA) or formaldehyde formation according to our previous reported method^17^.

##### (2) Glycine cleavage enabled by H_lip_ alone

The glycine cleavage reaction enabled by H_lip_ alone was monitored by determining the formation of NADH at 340 nm. The reaction mixture contained Tris-HCl (50 mM, pH 7.5), 0.5 mM THF, 20 mM DTT, 25 μM PLP, 5 mM NAD^+^, 40 μM FAD and 10 µM H_ox_. The reaction was initiated by adding 50 mM glycine.

#### The overall reaction of glycine synthesis

##### (1) Glycine synthesis catalyzed by GCS

The reaction mixture containing Tris-HCl (50 mM, pH 7.5), 0.5 mM THF, 20 mM DTT, 10 mM HCHO, 25 μM PLP, 5 mM NADH, 5 µM P-protein, 5 µM T-protein, 5 µM L-protein and 10 µM H_ox_. The reaction was initiated by adding 50 mM NH_4_HCO_3_ to the reaction mixture and carried out for 2 h at 37 °C. The amount of glycine formed was determined using HPLC. One unit of glycine synthesis activity was defined as the amount (in mg) of H-protein that catalyzed the formation of 1 nmol of glycine per minute.

##### (2) Glycine synthesis enabled by H_lip_ alone

The reaction mixture contained Tris-HCl (50 mM, pH 7.5), 0.5 mM THF (added with β-ME to prevent its oxidative degradation), 10 mM HCHO, 25 μM PLP, 50 mM NH_4_HCO_3_, 40 μM FAD and 5 mM NADH, and different concentration of H_ox_ (10-80 μM). Alternatively, 20 mM DTT can be used to replace FAD and NADH for the reduction of H_ox_ to H_red_. The reaction condition and enzyme activity calculation were the same as stated above in 2.3.2 (1).

#### Individual GCS reaction steps in the presence of H_lip_ with or without the corresponding enzymes

##### (1) Glycine dcarboxylation reaction catalyzed by P-protein

The reaction mixture contained Tris-HCl (50 mM, pH 7.5), 50 mM glycine, 50 µM H_ox_ and 25 µM PLP. 5 µM P-protein was added to the reaction mixture as the control group. The reaction was carried out for 2 h at 37 °C. The substrate H_ox_ and the product H_int_ were measured using HPLC.

##### (2) Aminomethyl transfer reaction catalyzed by T-protein

In this reaction of converting H_red_ to H_int_ through aminomethyl transfer, H_red_ required was generated by reducing H_ox_ with DTT (see 2.3.3 below), and 5,10-CH_2_-THF was derives from the condensation of HCHO and THF. Therefore, the reaction mixture contained Tris-HCl (50 mM, pH 7.5), 50 µM H_ox_, 0.5 mM THF, 20 mM DTT, 50 mM NH_4_Cl, and 10 mM HCHO. 5 µM T-protein was added to the reaction mixture as the control group. The reaction was carried out for 2 h at 37 °C. The substrate H_ox_ and the product H_int_ were measured by HPLC.

#### Electron transfer reaction between H_ox_ and H_<u>red</u>_ with or without the presence of L-protein

The interconversion of H_ox_ and H_red_ was performed according to a reported enzymatic assay using an excess amount of a reductant (8 mM TCEP or 20 mM DTT) for the reduction of the H-protein-bound lipoic acid^34, 35^, and then the produced H_red_ is re-oxidized by L-protein in the presence of NAD^+^. For the assay, the reaction mixture contained different types of buffer (50 mM, pH 7.5), 8 mM TCEP or DTT, 5 µM H_ox_ and 0.2 µM L-protein. In order to prove that H_ox_ can still undergo redox reaction without L-protein, 40 µM FAD is used instead of L-protein. The reactions were initiated by the addition of 5 mM NAD^+^. The rate of NADH formation was determined spectrophotometrically at 340 nm.

### Analytical methods using HPLC

H_ox_ and H_int_ proteins were analyzed based on the HPLC method previously developed in our lab^33^. The analysis was performed with a Inertsil WP300 C4 column (5 μm, 4.6×150 mm) and monitored at 280 nm using a diode array detector (DAD).The mobile phase consisted of acetonitrile (A) and 0.1 % trifluoroacetic acid aqueous solution (B). The volumn percentage of buffer B was varied as follows: linearly increased from 30 % to 50 % (0-13.4 min), sharply increased from 50 % to 90 % (13.4-13.41 min), held at 90 % (13.41-14.2 min), and then sharply decreased to 30 % (14.2-14.21 min), held at 30 % to 18 min. The flow rate was 1.0 mL·min^−1^.

Glycine concentration in the reaction mixture was determined by pre-column dansyl chloride derivatization. To this end, 40 µL of a reaction mixture was mixed with 160 µL of 0.2 M NaHCO_3_ and 200 µL of 5.4 mg · mL^−1^ dansyl chloride in acetonitrile. Derivatization occurred at 30 °C for 30 min. After the reaction, 600 µL of 0.12 M HCl was added to adjust the pH of the sample to weak acidic. After centrifuged at 10,000 rpm the supernatant was filtered with 0.22 µm membrane. The dansyl derivative of glycine was measured using HPLC (Shimadzu LC-2030C system) on a Shim-pack GIST C_18_ column (5 μm, 4.6×150 mm) at 30 °C, with a mobile phase composed of acetonitrile and 20 mM potassium phosphate buffer pH 6.0 (25:75 v/v) at a flow rate of 0.8 mL/min. The effluent was monitored at 254 nm using a diode array detector (DAD). The HPLC results of glycine were given in Supplementary materials Figure 2.

## Statistics and Reproducibility

Enzyme activities and reaction rates were measured by three independent experiments and averaged for report. Individual data points are added in the graphs, and error bars are defined by the standard deviation.

## Data availability

Major data generated and analyzed during this study are included in the article. The source data underlying the graphs and charts presented in the main figures are available as Supplementary Data. Other datasets generated and analyzed during the study are available from the corresponding author on reasonable request.

## Acknowledgments and funding

This work was financially supported by the Beijing Advanced Innovation Center for Soft Matter Science and Engineering, Beijing University of Chemical Technology.

## Author Contributions

Y.X. designed and performed the experiments, wrote the initial manuscript. Y.L. assisted in experiments and data analysis. H.Z. provided H-protein mutants and participated in data analysis. J.N. assisted in experiments and preparing the figures. J.R. involved in experimental design, data analysis and drafting the manuscript. W. W. involved in data analysis and revised most of the manuscript content. A.-P.Z. supervised the project, involved in experimental design, data analysis and discussion, reviewed and revised the paper.

